# Pirfenidone treatment attenuates fibrosis in autosomal dominant polycystic kidney disease

**DOI:** 10.1101/2025.08.25.672225

**Authors:** Viji Remadevi, Abeda Jamadar, Meekha M Varghese, Haichun Yang, Sumedha Gunewardena, Darren P Wallace, Reena Rao

## Abstract

Autosomal dominant polycystic kidney disease (ADPKD) is a leading genetic cause of kidney failure, marked by progressive cyst expansion, inflammation and fibrosis. Renal fibrosis, characterized by myofibroblast activation and excessive extracellular matrix (ECM) deposition is a central driver of disease progression in ADPKD, yet targeted anti-fibrotic therapies remain limited. Here, we evaluated the therapeutic potential of pirfenidone to suppress fibrosis and disease progression in ADPKD. To define the ECM in human ADPKD kidneys, we analyzed snRNA-seq data and found that fibroblasts are the principal source of fibrous and adhesive ECM in ADPKD kidneys, exhibiting higher ECM gene expression than normal controls. *In vitro,* primary culture human ADPKD renal myofibroblasts showed a similar profibrotic gene expression profile, and pirfenidone treatment suppressed ECM gene expression, cell proliferation, migration and contractility. In the Pkd1^RC/RC^ mouse model of ADPKD, pirfenidone treatment significantly reduced renal fibrosis, myofibroblast accumulation, ECM deposition, pro-fibrotic gene expression and associated cell signaling pathways. Pirfenidone also decreased kidney-to-body weight ratio and improved kidney function in Pkd1^RC/RC^ mice, without altering cyst burden. Collectively, these findings demonstrate that pirfenidone attenuates renal fibrosis and improves kidney function in ADPKD by targeting myofibroblast activation and ECM production, supporting a complementary therapeutic approach to cyst-directed therapies.

## INTRODUCTION

ADPKD is a common inherited monogenic disorder caused by mutations in the *PKD1* or *PKD2* genes. It is estimated that 42.6 per 100,000 persons in the United States and over 12.5 million people worldwide are afflicted with ADPKD (1, 2). Progressive cyst growth in the kidneys and liver, accompanied by interstitial fibrosis, inflammation and hypertrophy are hallmarks of ADPKD (3). Among these features, kidney fibrosis is a critical pathological component and a major contributor to declining kidney function and eventual progression to end-stage kidney disease (ESKD) (4–7).

Progressive kidney interstitial fibrosis leads to tissue scarring and disruption of normal kidney achitecture and loss of function, and contributes to systemic complications such as hypertension, metabolic dysregulation and electrolyte imbalance. In ADPKD, fibrosis is not merely a secondary consequence of cyst expansion but a key pathogenic driver of disease progression. ADPKD kidneys show excessive ECM accumulation, abnormal turnover and extensive remodeling alongwith activation of pro-fibrotic cell signaling pathways (4–6). This altered fibrotic microenvironment in turn promotes cyst growth. For instance, ECM accumulation contributes to cyst growth *via* integrin signalling (7, 8).

Myofibroblasts are the principal producers of ECM. They are highly contractile and migratory cells which express α-smooth muscle actin (αSMA). Myofibroblasts are capable of producing copious amounts of ECM and pro-fibrotic and pro-inflammatory factors that modify the interstitium in chronic kidney disease and chronic diseases of most solid organs (9, 10). While myofibroblasts are scarce in the normal kidney tubular microenvironment, they are abundant in ADPKD kidneys and often seen surrounding cysts. Our previous studies showed that in ADPKD kidneys, the cyst-lining epithelial cells drive myofibroblast activation and accumulation *via* paracrine signaling, establishing a pro-fibrotic pericystic microenvironment (11). Importantly, depletion of myofibroblasts in ADPKD mouse kidneys reduced both interstitial fibrosis and cyst growth, highlighting a unique pathogenic role for these cells in ADPKD (12).

Most ADPKD cases are diagnosed when cysts are already evident and kidney function begins to decline. Except in individuals with a known family history, ADPKD diagnosis mostly occurs because of comorbidities such as hypertension (13). Importantly, Tolvaptan is currently recommended only for use in patients with advanced cystic disease (Mayo-Irazabal classification 1D-1E) (14), by which stage, the kidneys are already fibrotic and continued ECM accumulation further compromises kidney function. Therefore, preventing or reversing kidney fibrosis remains a key therapeutic objective in ADPKD.

Pirfenidone (PFD) (5-methyl-1-phenyl-2-[1H]-pyridinone) is a pyridine derivative antifibrotic drug approved by the FDA for the treatment of idiopathic pulmonary fibrosis (IPF) (15). It shows antifibrotic, anti-inflammatory and antioxidant effects across preclinical and clinical models, including fibrotic lesions of the heart, liver, skin and pancreas (16). PFD is also kidney-protective in animal models of kidney injury including unilateral ureteral obstruction, nephrotoxicity, diabetic nephropathy and anti-glomerular basement membrane glomerulonephritis, mainly by targeting TGF-β signaling and reducing ECM deposition (17–19).

Given the central role of fibrosis in ADPKD progression, we investigated whether PFD could attenuate kidney fibrosis in the well-established Pkd1^RC/RC^ (RC/RC) mouse model of ADPKD. We also aimed to elucidate the underlying molecular mechanisms of its action, with a focus on the ECM, myofibroblast activity and key signaling pathways. Our findings demonstrate that PFD reduces myofibroblast activity, kidney fibrosis and cyst growth, and may be a potential therapeutic drug for the treatment of PKD.

## RESULTS

### PFD treatment reduced cell proliferation, migration and contractility of human ADPKD renal myofibroblasts

To test the effect of PFD in ADPKD, we first tested its effect on primary culture human ADPKD renal myofibroblasts (Fig. 1A). PFD treatment reduced ADPKD myofibroblast cell viability (Fig. 1B,C), and cell proliferation indicated by a 10-fold reduction in BrdU incorporation compared to vehicle-treatment (Fig. 1D). To examine the effect of PFD on migration, a scratch-wound healing assay was performed after mitomycin-pretreatment to block cell proliferation. Vehicle-treated ADPKD myofibroblasts closed the wound within 8 hours, while PFD-treated cells closed only ∼50% of the wound area over the same time period (Fig. 1E, F).

**Figure 1:**
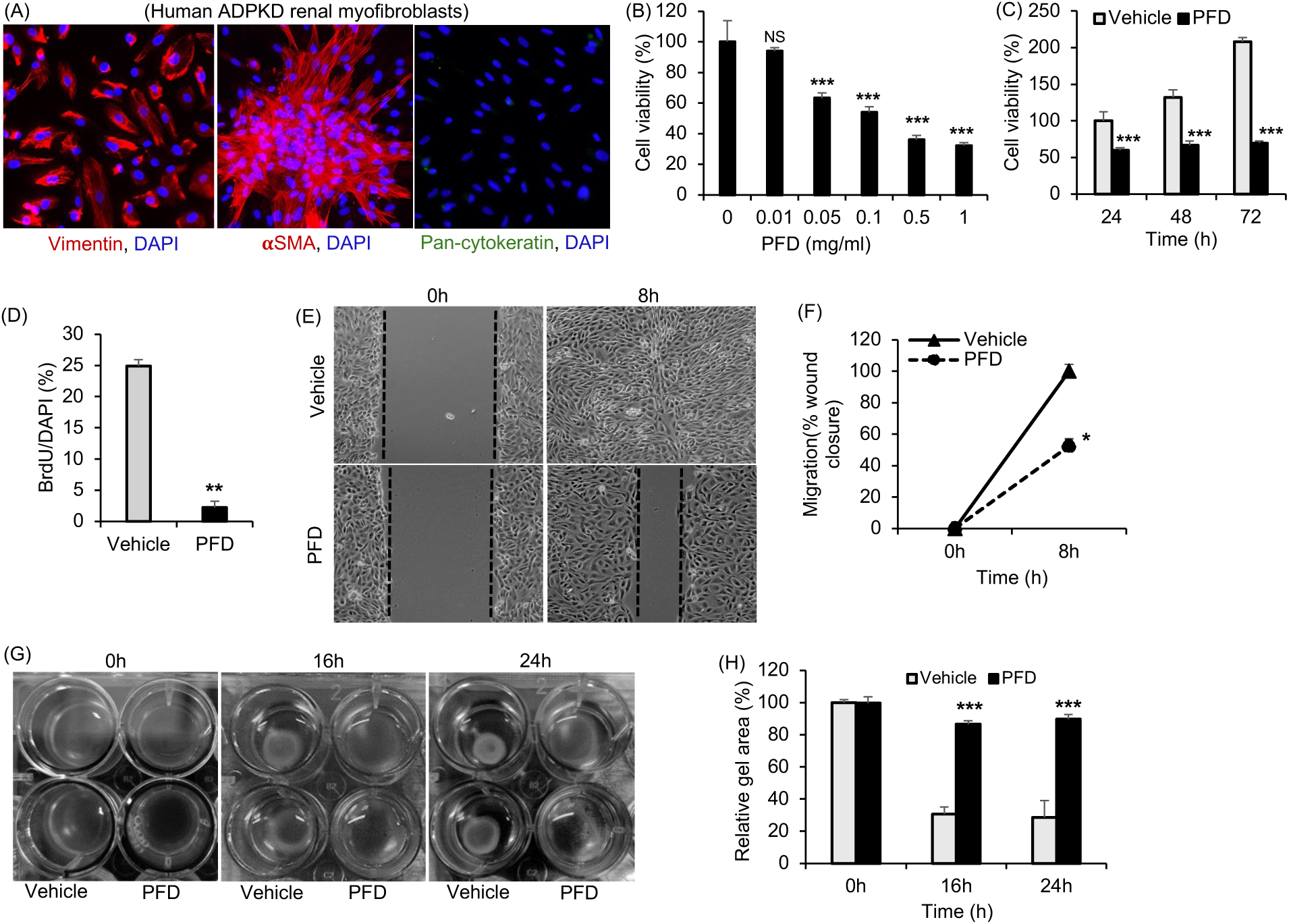
Effect of PFD on human ADPKD renal myofibroblasts. (A) Human ADPKD myofibroblasts stained for vimentin (denotes mesenchymal cells), αSMA (denotes myofibroblasts), and pan-cytokeratin (epithelail cell marker). (B) In human ADPKD myofibroblasts, cell viability assessed by MTT assay after PFD treatment for 24hrs. (C) Time course of cell viability in human myofibroblasts cells treated with 0.5mg/ml PFD. (D) Cell proliferation measured by BrdU assay after treatment with vehicle or PFD (0.5mg/ml for 24hrs). (E) Migration assay in cells treated with vehicle or PFD (0.5mg/ml), and (F) Quantification of migration. (G) Gel contractility assay in cells treated with vehicle or PFD (0.5mg/ml). (H) Relative area of the collagen lattice. *P<0.05, **P<0.01, ***P<0.001 by Unpaired t-test with Welch’s correction.

Myofibroblasts express αSMA, contributing to their contractile phenotype and driving tissue distortion and disease progression in solid organs (10). When ADPKD myofibroblasts were cultured in collagen gels, PFD treatment reduced gel contraction by ∼ 60% when compared to vehicle treatment at 16 and 24h (Fig. 1G, H).

### Myofibroblasts are a predominant source of ECM in human ADPKD kidneys

Myofibroblasts are the major producers of ECM in chronic kidney disease (10, 20, 21). However in ADPKD kidneys, the specific cell types that produce ECM, the ECM composition and their relative levels expressed by these cell types remain poorly defined. We analyzed the publicly available Kidney Interactive Transcriptomics (KIT) single-nuclei RNA sequencing (snRNA-seq) database comparing human normal kidneys (n=5) and ADPKD kidneys (n=8) (22). A cutoff of gene expression of >=1 was applied to identify genes actively transcribed in the cell clusters. Among the structural fibrous ECM proteins, we detected gene expression of 21 collagen subtypes, as well as elastin (*ELN*) and fibrillin-1 (*FBN1*) in kidney cell clusters (Fig. 2A*). COL4A* subtypes *COL4A3* and *COL4A4* exhibited the highest expression levels among all collagens, and were predominantly expressed in podocytes of both control and ADPKD kidneys (Fig. 2A). Compared to other kidney cell types, ADPKD fibroblast cluster was the major source of 13 collagen subtypes, including *COL1A1, COL1A2, COL3A1, COL4A1, COL4A2, COL5A1, COL5A2, COL6A2, COL6A3, COL12A1, COL14A1, COL15A1* and *COL16A1* (Fig. 2A). Notably, the expression of the above mentioned collagens, along with *COL8A1* and *FBN1* were elevated in the fibroblast cluster of ADPKD kidneys compared to normal control kidneys (Fig. 2A).

**Figure 2:**
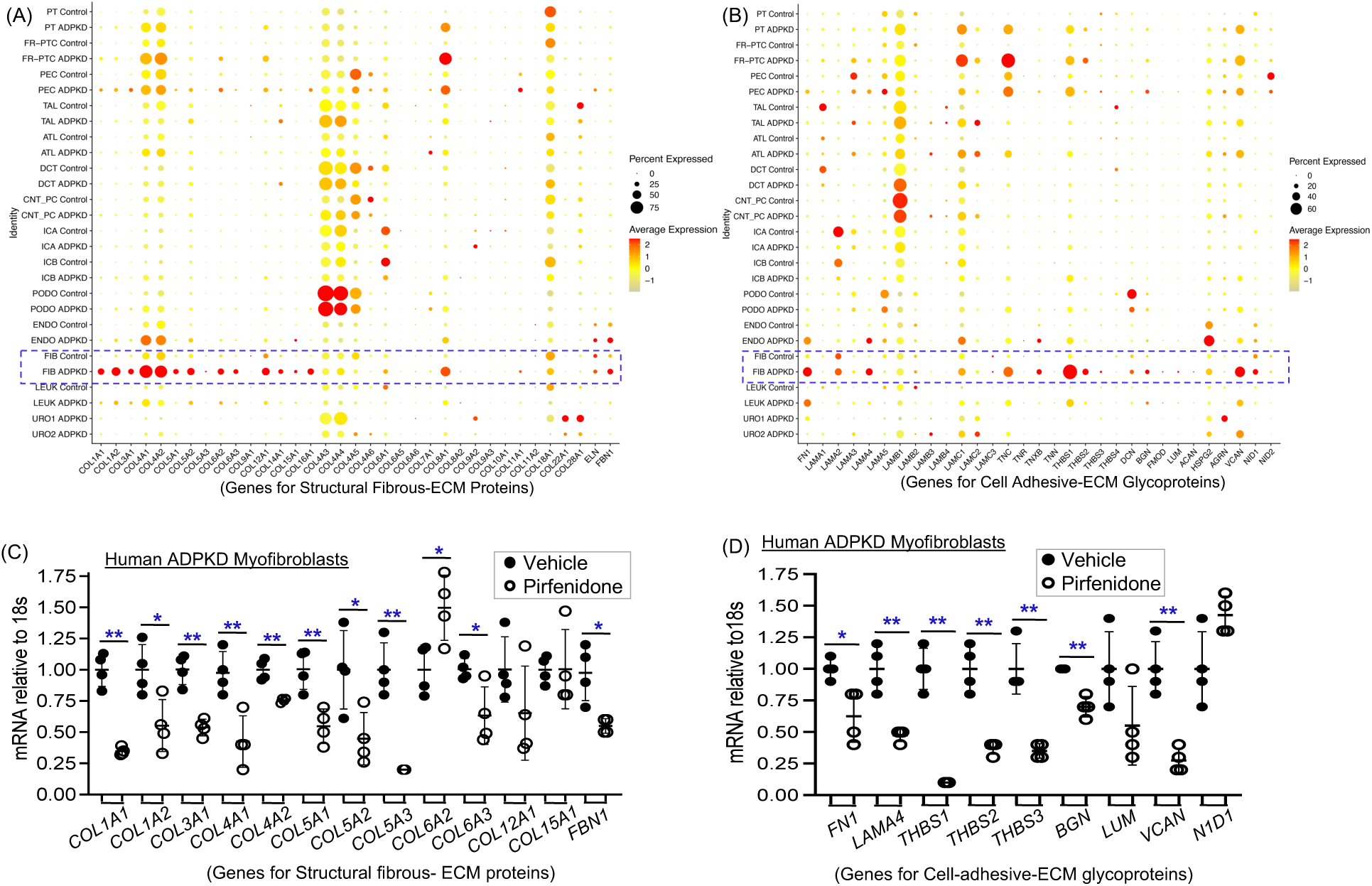
Effect of PFD on expression of structural fibrous-ECM proteins and cell adhesive ECM glycoproteins in human ADPKD renal myofibroblasts. (A) Dot plots representing the snRNA-seq-KIT dataset for human ADPKD (n=8) and human normal control (n=6) kidneys. Gene expression for structural fibrous ECM proteins and (B) cell adhesive ECM-glycoproteins are shown. The size (diameter) of each dot reflects the percentage of cells expressing the gene, while the color intensity indicates the average expression level compared to all cell types. (C) Effect of PFD (0.5mg/ml for 24h) or vehicle treatment on mRNA levels of structural fibrous ECM proteins and (D) cell adhesive ECM-glycoproteins in human ADPKD myofibroblasts. *P<0.05, **P<0.01 by Unpaired t-test with Welch’s correction.

Of the 18 genes of cell adhesive ECM glycoproteins expressed in ADPKD kidneys (Fig. 2B), ADPKD fibroblast cluters were the major expressors of 9 genes including fibronectin (*FN1*), laminin subunit α4 (*LAMA4*), tenascin-XB (*TNXB*), thrombospondin-1 (*THBS1*), *THBS2*, biglycan (*BGN*), versican (*VCAN*), and nidogen-1 *(NID1*), compared to other kidney cell types. The above mentioned glycoproteins, along with tenascin-C (*TNC*), *LAMC1*, *LAMC2*, and perlecan (*HSPG2*) showed significantly higher expression in ADPKD fibroblasts relative to normal control kidney fibroblasts (Fig. 2B).

### PFD treatment reduced ECM in human ADPKD renal myofibroblasts

To validate the findings from the snRNA-seq analysis, we examined gene expression of ECM proteins in primary culture human ADPKD myofibroblasts. Consistent with SnRNA-seq data, most structural fibrous ECM proteins and cell-adhesive ECM glycoproteins identified in ADPKD fibroblasts were also detected in primary culture ADPKD myofibroblasts (Supplemental Fig. 1A, 1B). Notably, PFD treatment significantly reduced gene expression of most ECM proteins except *COL12A1, COL15A1, LUM* and *NID1* (Fig. 2C,D). *COL6A2* was an exception, which showed significantly increased levels in PFD treated cells when compared to vehicle treatment (Fig. 2C).

### PFD treatment reduced renal fibrosis in RC/RC mice

To determine the effect of PFD on renal fibrosis in ADPKD, male RC/RC mice were treated with vehicle or PFD (200mg/kg, twice-a-day by oral gavage), from 4 months of age to 6 months of age and sacrificed. PFD treatment significantly reduced gene expression of multiple collagens, including *Col1a1, Col1a2, Col3a1, Col4a2, Col5a1, Col5a2, Col5a3, Col6a1, Col6a2, Col8a1, Col12a1,* and *Col18a1* (Fig. 3A). While *Col4a1* and *Col15a1* were also detected, their expression levels were not significantly altered (Supplemental Fig 2). Additionally, the expression of *Fn1*, *Lama4, Thbs1* and *Thbs2* were significantly reduced in PFD-treated RC/RC mouse kidneys compared to vehicle-treated controls (Fig. 3B).

**Figure 3:**
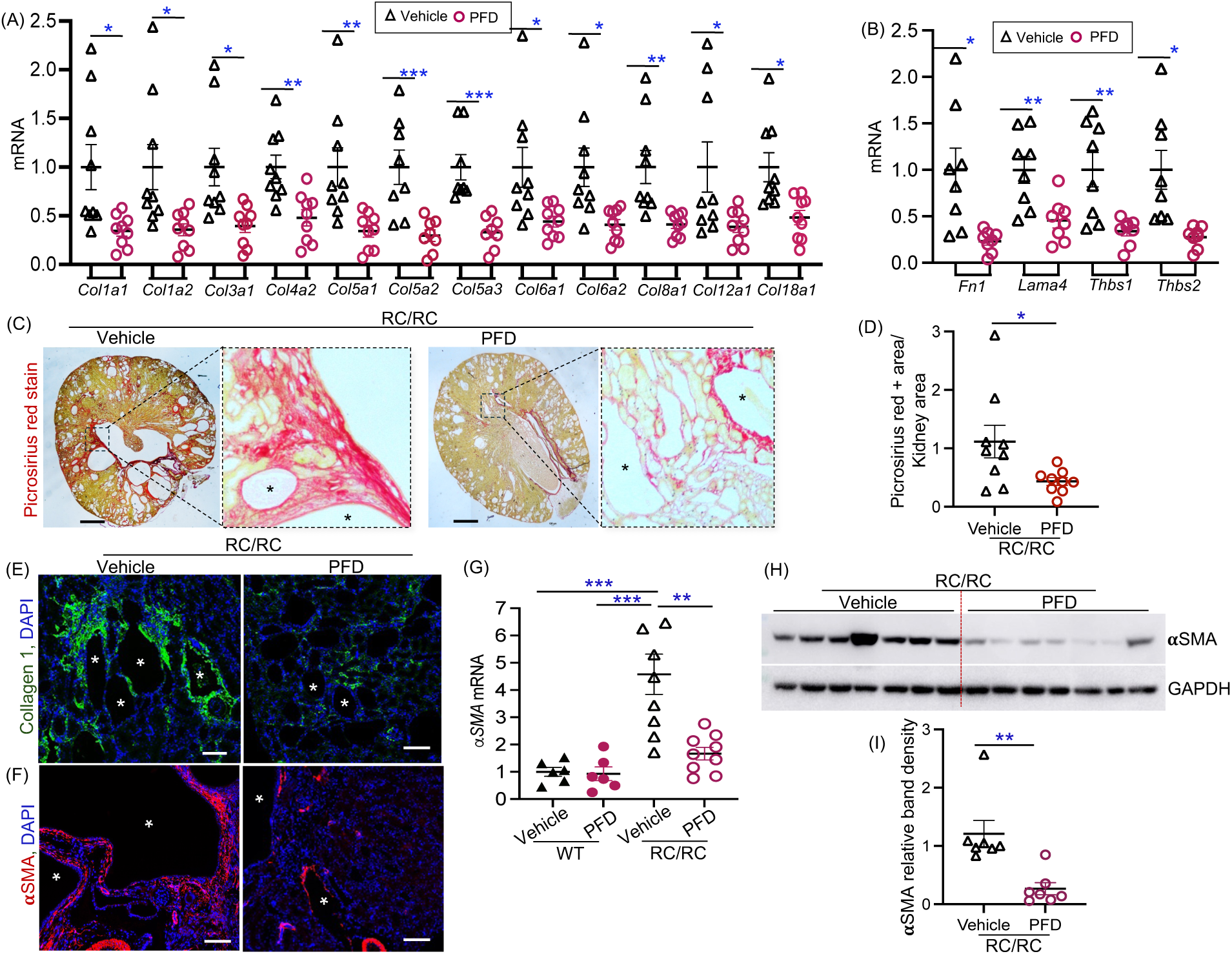
Effect of PFD on fibrosis in RC/RC mouse kidneys: (A) Kidney mRNA levels of structural fibrous ECM proteins, and (B) cell adhesive-ECM glycoproteins. (C) Sirius Red staining of mouse kidney tissue (Scale bar 1mm), and (D) Quantitation of Sirius Red staining. (E) Kidney tissue immunostained for Collagen 1A (green) and DAPI (nuclei, blue) (Scale bar 100µm), and (F) αSMA (red) (Scale bar 100µm). (G) αSMA mRNA levels relative to 18S in kidney tissue. (H) Western blot analysis of the whole kidney tissue lysate, and (I) Densitometry of immunoblots. *P<0.05, **P<0.01, ***P<0.001 by ordinary one-way ANOVA in G, and Unpaired t-test with Welch’s correction in all others. Star symbols in figures C, E and F represent cysts.

PFD treated RC/RC mouse kidneys showed significantly reduced kidney fibrosis when compared to vehicle treated RC/RC mice, as demonstrated by reduced Picrosirius Red staining and quantitation (Fig. 3C, D). PFD-treated RC/RC kidneys also showed reduced collagen-1 immunostaining compared to vehicle treated RC/RC kidneys (Fig. 3E).

We next examined the effect of PFD treatment on the expression of αSMA, a well-established myofibroblast marker. In vehicle treated RC/RC kidneys, immunostaining revealed αSMA expressing cells surrounding the cysts. PFD treatment markedly reduced αSMA expression in the RC/RC kidney (Fig. 3F), indicating a reduction in myofibroblasts. Quantitative analysis confirmed a significant decrease in both *Acta2* (αSMA) mRNA levels (Fig. 3G) and αSMA protein levels (Fig. 3H,I) in PFD-treated mice compared to vehicle-treated controls.

### PFD attenuates fibroblast-enriched ECM remodeling and pro-fibrotic signaling in ADPKD

To examine the mechanism underlying the anti-fibrotic effect of PFD in ADPKD kidneys, we first examined the expression of ECM remodeling matrix metalloproteinases (MMPs), their tissue inhibitors (TIMPs) and A Disintegrin and Metalloproteinases (ADAMs). In addition, we examined matricellular proteins which play critical roles in regulating cell-matrix signaling and myofibroblast activation. Analysis of KIT snRNA-seq database showed that gene expression of *TIMP1, TIMP2, TIMP3, ADAM12* and *ADAM17* genes were higher in fibroblasts compared to other cell types in human ADPKD kidneys (Fig. 4A). The above mentioned genes, and *MMP7, ADAM10* and *ADAMTS1* exhibited higher expression levels in fibroblast clusters of ADPKD kidneys compared to normal control kidneys (Fig. 4A). Matricellular proteins, including osteopontin (*SPP1*), cellular communication network factors (*CCN1* and *CCN2*), osteonectin (*SPARC*), fibulin-1 (*FBLN1*), *FBLN5* and *SERPINE1* exhibited higher expression in fibroblast clusters from ADPKD kidneys compared to normal control kidneys (Fig. 4B).

**Figure 4:**
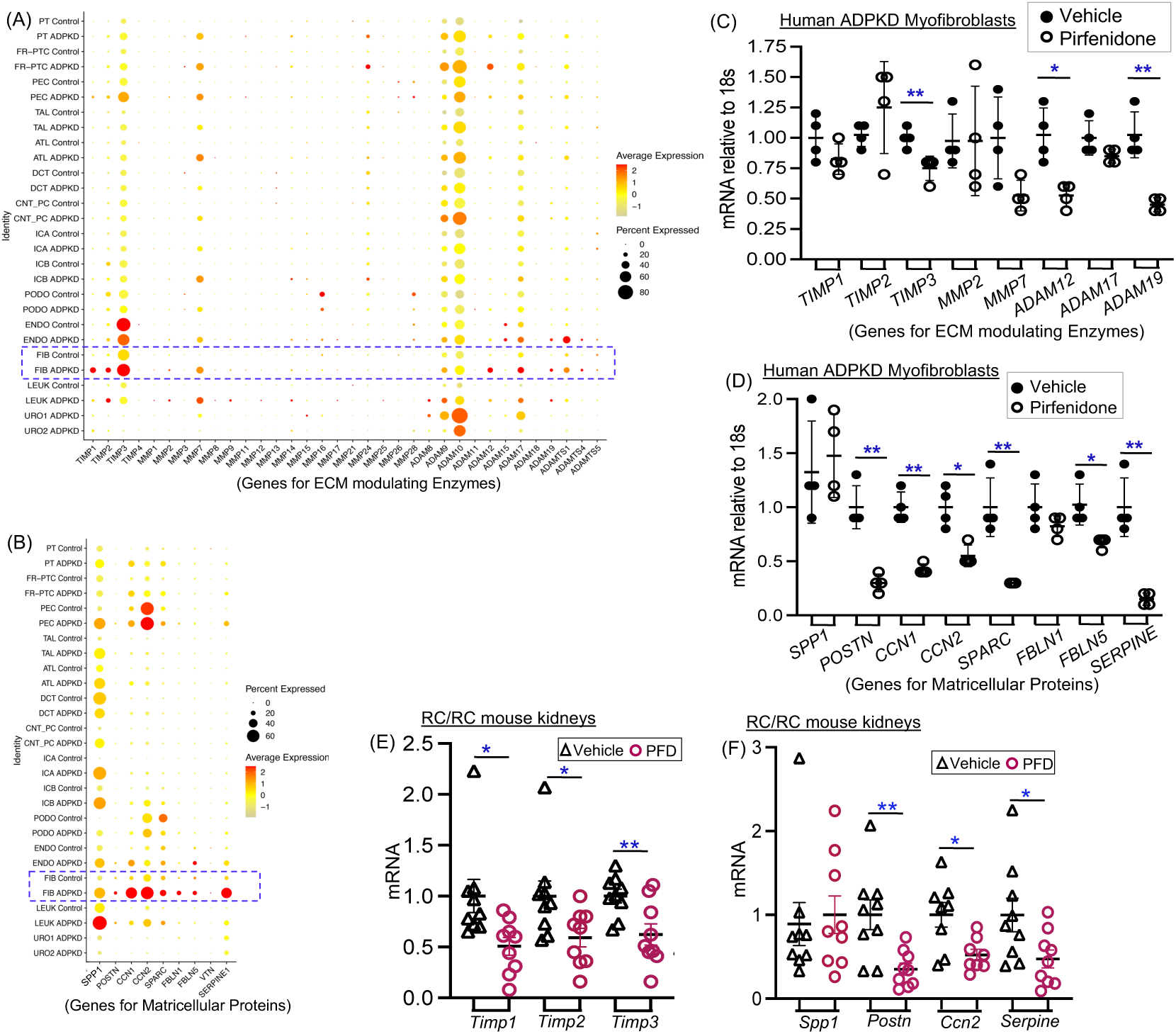
Effect of PFD on ECM modulating enzymes and matricellular proteins in ADPKD. (A) Dot plots representing the snRNA-seq KIT dataset as in Fig. 2 show gene expression of ECM-modulating enzymes, and (B) matricellular proteins. (C) In human ADPKD renal myofibroblasts, the effect of PFD (0.5mg/ml for 24h) or vehicle treatment on mRNA levels of ECM modulating enzymes, and (D) matricellular proteins. (E) In RC/RC mouse kidney tissues, mRNA levels of ECM modulating enzymes and (F) matricellular proteins. *P<0.05, **P<0.01 by Unpaired t-test with Welch’s correction.

In primary culture human ADPKD myofibroblasts, PFD treatment significantly reduced gene expression of *TIMP3, ADAM12 and ADAM19* (Fig. 4C); and POSTN, *CCN1, CCN2, SPARC, FBLN5* and *SERPINE1* (Fig. 4D). In RC/RC mouse kidneys, PFD treatment reduced *Timp1, Timp2* and *Timp3* expression (Fig. 4E); and reduced *Postn1, Ccn2* and *Serpine-1* (Fig. 4F) compared to vehicle treatment. *Mmp2, Mmp7* and *Mmp9* showed no significant differences in RC/RC kidneys (Supplemental Fig. 3).

Based on the above *in vitro, in vivo* and snRNA-seq KIT data TIMP3, CCN2 and TSP1 are factors expressed in human and ADPKD kidneys, and are significantly reduced by PFD treatment. Hence, we focused on common pro-fibrotic signaling pathways regulated by these factors, particularly TGFβ/SMAD3, ERK1/2, YAP, AKT and β-catenin. PFD-treatment significantly decreased SMAD3 and AKT activity indicated by reduced pSMAD3/SMAD3 and pAKT/AKT ratios; and also reduced YAP and β-catenin levels in RC/RC mouse kidneys (Fig. 5A,B,C,D). ERK1/2 activity showed no significant difference between PFD and vehicle treated RC/RC mouse kidneys (Fig. 5C, D).

**Figure 5:**
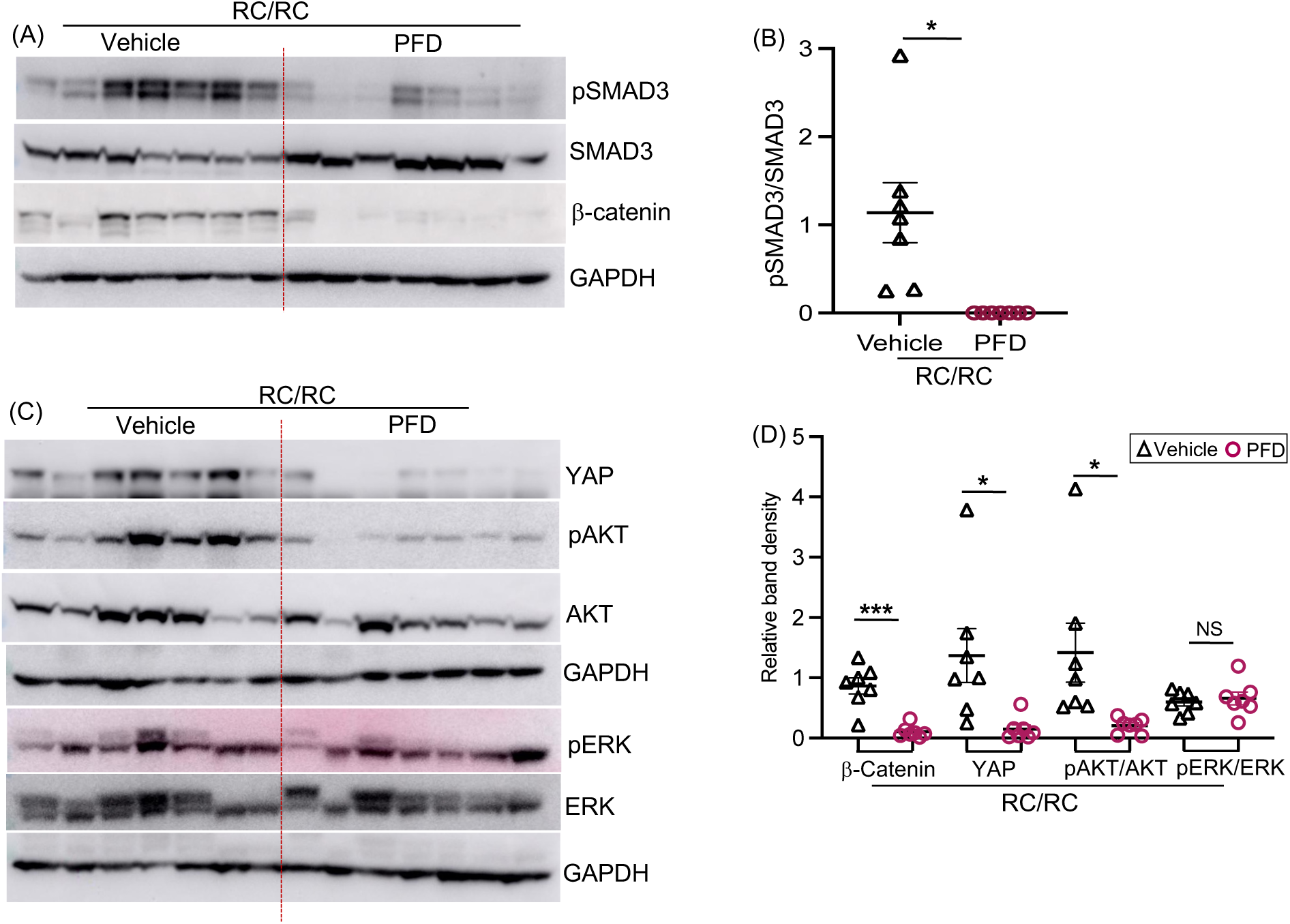
Effect of PFD on profibrotic cell signaling. (A) Western blot analysis on whole kidney tissue lysate for, (A) pSMAD3/SMAD3 and β catenin and (B) Quantification of band density for pSMAD3/SMAD3, (C) YAP, pAKT/AKT and pERK/ERK and (D) Quantification of band density. *P<0.05, **P<0.01, ***P<0.001 by Unpaired t-test with Welch’s correction.

Supporting our *in vivo* findings, TGFβ treatment of cultured NRK-49F rat kidney fibroblasts significantly induced *Acta2* (αSMA) mRNA (Supplemental Fig. 4A) and protein expression (Supplemental Fig. 4B, C) which were completely abolished when treated with PFD (Supplemental Fig. 4A,B,C). Collectively, these findings demonstrate that PFD attenuates fibroblast-driven ECM remodeling and matricellular signaling programs, leading to suppression of key pro-fibrotic pathways in ADPKD kidneys.

### PFD treatment reduced cyst growth in RC/RC mouse kidneys

PFD treatment significantly reduced kidney-to-body weight ratio (Fig. 6A) and kidney weight (Fig. 6B) in RC/RC mice, without significantly affecting overall body weight compared to vehicle-treated controls (Supplemental Fig. 5). PFD treatment did not significantly alter cystic index (Fig. 6C) or cyst number (Fig. 6D) in RC/RC kidneys. However, PFD treated RC/RC mice showed significantly reduced blood urea nitrogen (BUN) levels (Fig. 6E) and more intact kidney parenchyma with less interstitial expansion compared to vehicle treated RC/RC mice (Fig. 6F,G). PFD treatment did not significantly alter mRNA expression of injury markers such as Kim-1 and Ngal (Supplemental Fig. 6A, B). In wildtype mice mice, PFD treatment showed no effect on kidney morphology (Fig. 6F, G), kidney-to-body weight ratio (Fig. 6A), kidney weight (Fig. 6B), or BUN (Fig. 6E) compared to vehicle treatment.

**Figure 6:**
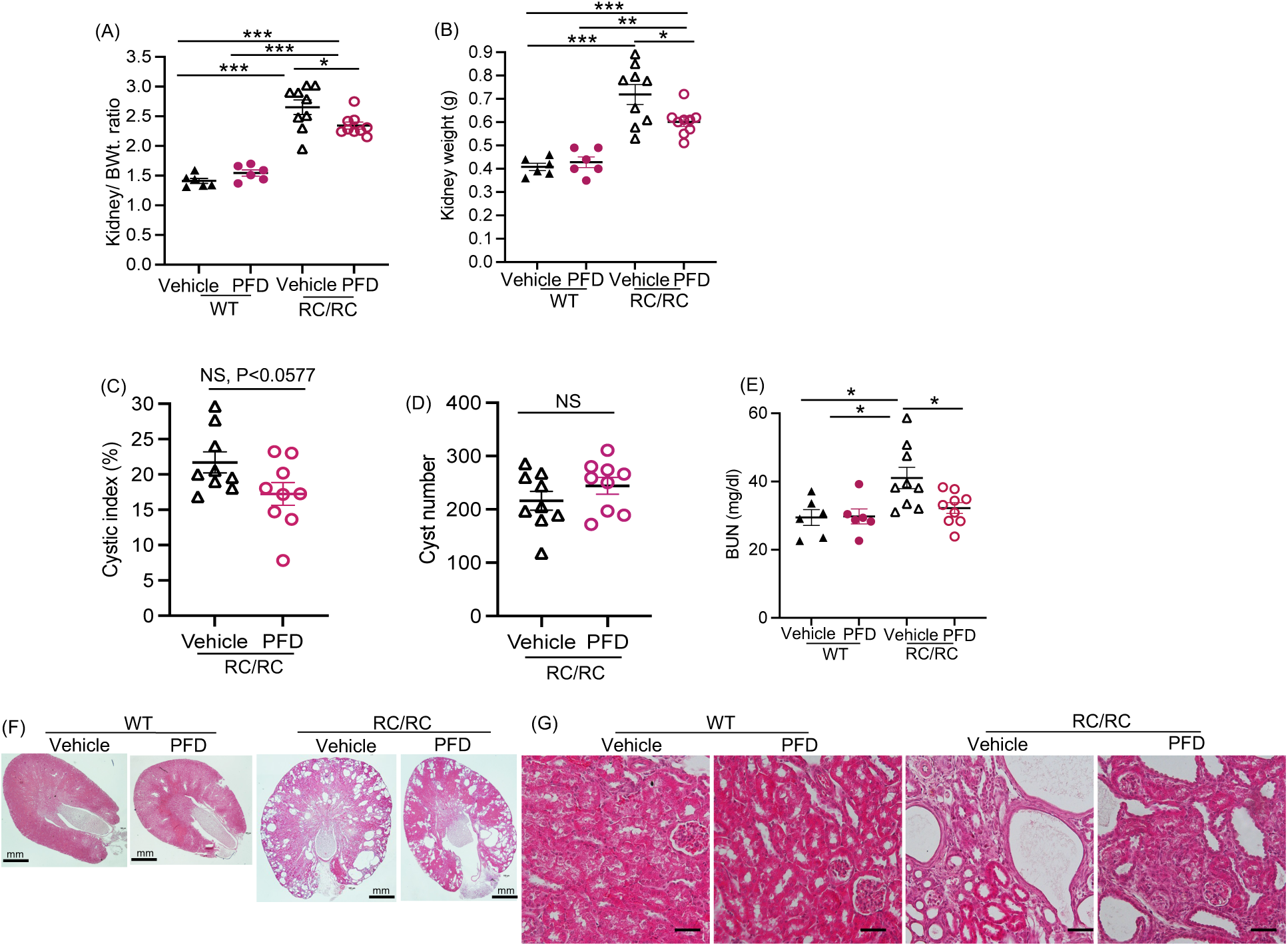
Effect of PFD on cyst growth in RC/RC mice: (A) Kidneys to body weight ratios, (B) Total kidney weight, (C) Cystic index, (D) Cyst number and (E) BUN. (F) Representative images of H&E staining (Scale bar 1mm), (G) Magnified images (Scale bar 100µm). *P<0.05, **P<0.01, ***P<0.001 by Unpaired t-test with Welch’s correction for C and D; and by ordinary one-way ANOVA with Tukey’s multiple comparison test in A, B and E.

## DISCUSSION

This study shows that PFD effectively attenuates pro-fibrotic processes in human ADPKD myofibroblasts and kidney fibrosis in the RC/RC mouse model of ADPKD. We found that myofibroblasts are a major source of ECM in human ADPKD kidneys, and that PFD treatment inhibits ECM gene expression, cell proliferation, migration and contractility of primary human ADPKD kidney myofibroblasts *in vitro*. In RC/RC mouse kidneys, PFD treatment reduced myofibroblast abundance, ECM, pro-fibrotic cell signaling pathways and kidney to body weight ratio, and improved kidney function compared to vehicle-treated controls. Together, these findings demonstrate that PFD attenuates fibrosis in ADPKD, in part by targeting myofibroblast-driven pathways.

We evaluated the effect of PFD on fibrosis in a mouse model of ADPKD as well as primary culture human ADPKD kidney myofibroblasts, to better reflect the human disease context. Given the limited characterization of ECM composition in ADPKD kidneys, we analyzed the publically available KIT snRNA-seq database, comparing normal and ADPKD human kidneys (22). We found that ADPKD fibroblasts are the major producers of multiple structural fibrous-ECM proteins, cell-adhesive ECM glycoproteins, ECM modulating enzymes and matricellular proteins when compared to other cell types. Moreover, compared to control normal fibroblasts, ADPKD fibroblasts exhibited elevated expression of numerous ECM and ECM-related genes, many of which are implicated in the pathogenesis of kidney diseases. For instance, type I (*COL1A1, COL1A2*) and type III (*COL3A1*) collagens are known to promote interstitial fibrosis, ECM stiffening, myofibroblast activation and vascular remodeling in the kidneys (23). Moreover, Type IV collagens (*COL4A1, COL4A2*), which are normally integral to basement membranes can become pathogenic when mutated or overexpressed (24). Type V (*COL5A1, COL5A2, COL5A3*) and type VI (*COL6A2, COL6A3*) collagens are also implicated in ECM remodeling and progression of fibrosis (25). Similarly, periostin, fibrillin-1, fibronectin, thrombospondin-1, biglycan and versican contribute to fibrosis by modulating TGFβ signaling and matrix organization (26–29).

In the current study, the ECM expression profile from KIT snRNA-seq data was validated in primary cultures of human ADPKD kidney myofibroblasts. Importantly, PFD treatment significantly reduced the expression of a broad panel of ECM-related genes in these cells. While most ECM and profibrotic genes were consistently downregulated by PFD in both human ADPKD myofibroblasts and RC/RC mouse kidneys, COL6A2 displayed a divergent response, showing increased expression in human ADPKD myofibroblasts but not in RC/RC mouse kidneys. This discrepancy may reflect compensatory mechanisms or cell-type specific responses to antifibrotic therapy and merits further investigation. Beyond its effects on ECM gene expression, PFD also significantly suppressed fundamental cellular behaviors of activated myofibroblasts including cell proliferation, migration and matrix contraction. Collectively, these findings highlight the potential of PFD to blunt both molecular and cellular drivers of fibrosis in ADPKD.

Consistent with the *in vitro* findings, PFD treatment significantly reduced the kidney myofibroblast population and ECM in RC/RC mouse kidneys. Mechanistically, our studies indicate that PFD treatment significantly suppressed TGFβ/SMAD3 signaling, which aligns with previous reports identifying inhibition of TGFβ signaling as a primary mechanism underlying PFD’s antifibrotic action (18). Additionally, PFD downregulated ECM remodeling enzymes and pro-fibrotic matricellular signaling pathways including AKT, β-catenin and YAP cell signaling. Thus, PFD holds potential as a therapeutic strategy to mitigate kidney fibrosis in ADPKD.

ECM remodeling and matrix stiffening are known to promote cyst growth in ADPKD through aberrant integrin and matricellular signaling (7, 30–32). Similarly, our prior studies show that depletion of αSMA positive myofibroblasts in RC/RC mice reduces both renal fibrosis and cyst growth (12). Moreover, human ADPKD renal myofibroblast conditioned media promotes cyst epithelial proliferation *in vitro*, highlighting their contribution to cystogenesis (12). Collectively, these findings indicate that modulation of the fibrotic microenvironment disrupts key paracrine and mechanotransductive cues that sustain cyst expansion in ADPKD. In the current study, systemic administration of PFD suppressed pro-fibrotic signaling pathways, and reduced myofibroblast abundance and ECM accumulation in RC/RC kidneys. PFD treatment also significantly decreased the kidney to body weight ratio and improved kidney function in RC/RC mice, indicating an overall attenuation of kidney enlargement. Although PFD treated RC/RC kidneys showed a trend towards reduced cyst index, this did not reach statistical significance compared to vehicle treated controls. Taken together, these findings support a model in which targeting fibrosis using PFD modifies the disease-permissive microenvironment and functionally restrains ADPKD progression, although the impact of PFD on cyst burden may require longer treatment duration or earlier intervention.

Notably, PFD’s antifibrotic effects occurred without evidence of nephrotoxicity, as kidney injury markers (*KIM1* and *NGAL*) remained stable and BUN levels improved. PFD showed favorable pharmacokinetics in humans including rapid absorption (Tmax: 0.33-1h) with a terminal half-life of 2-2.5 hours, low inter-individual variability, and minimal accumulation with multiple dosing (33). Phase III trials and post-marketing study data for IPF also confirm PFD’s safety and tolerability (34, 35). The common side effects included nausea, fatigue, weight loss and rash, with dose-dependent toxicities (GI, liver, skin) manageable through dose adjustment (36). In a previous study, we showed that nintedanib, a triple tyrosine kinase inhibitor that is also an FDA approved anti-fibrotic drug for IPF, reduced kidney fibrosis and cyst growth, and preserved kidney function in ADPKD (37). Although nintedanib has a broader range of action because of its kinase inhibition, it has more gastrointestinal side effects compared to PFD (38).

Given that current treatments for ADPKD, such as tolvaptan, primarily target cyst growth pathways, PFD may provide a complementary approach by addressing fibrotic remodeling. Future studies could assess the long-term effects of combination therapy on kidney function decline, cyst growth and fibrosis.

## MATERIALS AND METHODS

### In vivo study

#### Sex as a biological variable

Only male mice were used in this study. A careful anlaysis of 6-months old RC/RC mice on BALB/c background from our inbred colony showed no significant difference in kidney to body weight ratio or cystic index between male and female mice (Supplemental Fig 7A,B). However, we found significant increase in Collagen 1a1 and Collagen 5a1 mRNA levels in male mice compared to female mice, but no significant difference in Collagen 3a1 or αSMA levels were observed (Supplemental Fig 7C). Although a previous study showed increased disease progression in female RC/RC mice on C57BL/6J background (39), a later study showed no such sexual dimorphism in the RC/RC mouse model, especially on the BALB/c background (40). Because the molecular pathways examined are not sex-specific and are broadly implicated in ADPKD pathogenesis, the findings in the male mice are expected to be relevant to both sexes; however, future studies in females are warranted to confirm generalizability.

#### ADPKD mouse model

Pkd1^RC/RC^ (RC/RC) mouse (41, 42) is a slow progressing, adult, orthologous model of ADPKD carrying a temperature sensitive folding hypomorphic mutation (R3277C) in the Pkd1 gene. Mice are on pure BALB/c background, and inbred. Male WT and RC/RC mouse littermates were treated with vehicle (5% DMSO+1% Hydroxymethyl cellulose) or pirfenidone (#HY-B0673, MedChemExpress, USA) (200mg/kg BWt., twice daily) by oral gavage between 8-9 AM and 4-5 PM, 6 days a week, from 4 to 6 months of age and sacrificed at 6 months of age. Power analysis conducted based our previous study (37) determined that using 8 mice will provide 80% statistical power to detect a 23% reduction in kidney to body weight ratio in RC/RC mice treated with drug compared to controls (significance level (α) 0.05). Mice from each litter were assigned to study groups as they reached 4 months of age. No randomisation was used. Investigators performing the study and analyses were blinded to the identity of the mice. Mice were housed in a temperature controlled environment in a 12 hr light /12hr dark cycle. Experimental mice were placed on the same rack. All mice were sacrificed between 11:30 AM and 12:30 PM. Blood was collected and plasma isolated. Kidneys were weighed and flash frozen, or fixed in 4% paraformaldehyde. No adverse effects were observed and no mice or datapoints were excluded. Studies were approved by University of Kansas IACUC committee and ARRIVE guidelines (43) were followed.

#### H&E staining and quantification of cysts

H&E staining was performed on 5μm thick kidney tissue sections and imaged using Nikon 80i upright microscope, Tokyo, Japan. Cyst number and cystic index (cyst area / total area per kidney tissue section) were quantified using ImageJ, Madison WI, USA by an observer blinded to the sample’s identity (44).

#### Picros Sirius Red staining and quantification of tissue fibrosis

Formalin-fixed, paraffin-embedded mouse kidney sections were stained with Picrosirius Red (45) according to manufacturer’s instructions (ScyTek Laboratories, #PSR-2, Logan, UT, USA) to detect collagen fibers. Kidney fibrosis was quantitatively evaluated using polarized light microscopy in combination with image analysis software (QuPath). Collagen accumulation was determined by calculating the proportion of birefringent staining relative to the total renal tissue area.

#### Blood-urea nitrogen levels (BUN)

Plasma BUN was measured using commercially available QuantiChrom Urea Assay Kit (#DIUR-100, BioAssay Systems, Hayward, CA, USA) (46).

#### Western blot

Kidneys were homogenized in SDS Laemmli buffer and run in 10% SDS-polyacrylamide agarose electrophoresis gels (47). Primary antibodies for αSMA (#ab5694; Abcam, Cambridge, MA,USA), pERK1/2 (#9102S), ERK1/2 (#9101S), β-catenin (#9582S), pAKT (#4060S), AKT (#4691T), pSMAD3 (#9520T) and SMAD3 (#9513S) from Cell signaling, Danvers, MA, USA; YAP (#SC-101199) and GAPDH (#SC-32233) from Santa Cruz Biotechnology, Inc, Dallas, TX, USA, and anti-mouse (#P0447) and anti-rabbit (#P0448) secondary antibodies from Dako (Santa Clara, CA, USA) were used. Immunoreactive proteins were detected using ECL reagent (Amersham, GE Healthcare, Buckinghamshire, UK).

#### Immunofluorescence staining

Fixed and paraffin-embedded tissues sections were processed as previously described (48). Primary antibodies αSMA (#ab5694) from Abcam (Cambridge, MA, USA) and Collagen Type-1a (#203002 from MD Bioproducts, Oakdale, MN) were applied followed by incubation with secondary antibodies anti-Rabbit IgG Alexa fluor^@^ 488 and anti-Goat IgG Alexa fluor^@^ 594 from Invitrogen (Carlsbad, CA, USA). After incubation, tissue sections were washed, stained with DAPI, and mounted using Flour-G (Invitrogen, Carlsbad, CA, USA). Images were captured using a Nikon 90i upright microscope (Tokyo, Japan).

#### Quantitative real time PCR

RNA was isolated from whole kidney lysate using trizol method and cDNA prepared with High-capacity cDNA reverse transcription kit (# 4368814, Applied Biosystems, Foster City, CA, USA). SYBR Green PCR master mix (# A25742, Applied Biosystems, Foster City, CA, USA) was used for QRT-PCR (49). Supplemental Table-1 shows the primer sequences.

### In vitro studies

#### Primary culture human ADPKD myofibroblasts

Cells were isolated from human ADPKD kidney tissues obtained from the University of Kansas PKD Biomarkers and Biomedical core (37, 50). Cells were used in their first passage and grown in DMEM:F12 media with 10% FBS and 1% Pen/Strep. Mycoplasma contamination was monitored using DAPI staining and confirmed using PCR based mycoplasma testing kit.

#### NRK-49F Rat kidney fibroblasts

(#CRL-1570, ATCC^©^, Manassas, VA, USA) were grown in DMEM medium with 5% FBS and 1% Pen/Strep.

#### Cell viability and cell proliferation assays

MTT assay was conducted to assess cell viability (42). To measure cell proliferation by BrdU incorporation assay (42), human ADPKD kidney myofibroblasts grown on coverslips were serum starved overnight followed by release into media containing 0.2% FBS with PFD or vehicle. After 24h, cells were incubated with 3μg/ml BrdU (#10280879001, Millipore Sigma, Burlington, MA) for 3h, and cell proliferation was measured as BRDU/DAPI expressed as %.

#### Migration assay

Confluent monolayers of human ADPKD kidney myofibroblasts were treated with 5μg/ml mitomycin c (# M7949; Sigma Aldrich; MO; USA) for 2h; washed twice with PBS and a single scratch (wound) was created manually across the monolayer using a sterile pipette tip, and cell debris washed off using PBS. The cells were treated with vehicle or PFD. The wound was imaged at regular intervals using Phase contrast microscope until the wounds in any one study group closed 100%, and wound closure was quantified (51).

#### Gel contractility assay

Human ADPKD myofibroblasts were trypisinized and resuspended in complete medium, and mixed with rat-tail collagen type-1 (pH 7.4) (# 354236, Corning, Glendale, Arizona). 500μl of collagen/cell mixture containing 1.5 X 10^5^ cells was dispensed into 24-well cell culture plates coated with 0.2% BSA. The mixture was incubated to polymerize at 37°C in a CO_2_ incubator for 1hr. After polymerization, the gels were gently detached from the sides and were incubated with serum free medium. The gels were imaged at different time points and the gel area was calculated using image-J software.

#### Single-nucleus RNA sequencing

The snRNA-Seq data was obtained from the Kidney Interactive Transcriptomics data base by Muto, Y. et al.(22) (GSE185948) https://humphreyslab.com/SingleCell/search.php). The bioinformatics analysis of this data was performed in line with the description given by the authors in their Methods section. We recreated the exact R environment used by the authors in order to obtain results comparable to their’s for genes of our interest presented in the dot-plots.

#### Statistics

Values were expressed as mean ± standard error (SEM) for *in vivo* studies, and mean ± standard deviation for *in vitro* studies. The data were analyzed by two-tailed unpaired *t*-test with Welch’s correction, or one-way ANOVA followed by Tukey’s multiple comparisons test. P≤0.05 was considered statistically significant. Analysis was done using GraphPad Prism software Version 10 (GraphPad Software, Inc., La Jolla, CA, USA).

#### Study approval

Mouse studies were approved by University of Kansas IACUC committee and ARRIVE guidelines (43) were followed.

## Supporting information

Supplemental figure and Table 1 and will be used for the link to the file preprint site

## Data availability

All *in vivo* studies using mouse models were performed and tissues analyzed at the University of Kansas Medical Center. All data that was generated are provided in the figures in the main text or in Supplementary Material.

All requests for data will be processed based on institutional policies for noncommercial research purposes and shared. Data sharing could require a data transfer agreement as determined by the University of Kansas Medical Center’s legal department.

## Author contribution

RR conceptualized and designed studies, analyzed results, and wrote the paper. VR performed experiments, analyzed results and wrote parts of the paper. AJ, MMS and HY performed some studies and analyzed results. SG performed bioinformatics analysis and generated the dot plots from the publically available Kidney Interactive Transcriptomics database. DPW conceptualized some studies and reviewed the paper. All authors read, edited and approved the paper.

## Acknowledgement

We thank Dr. Peter Harris and Dr. Katharina Hopp for providing *Pkd1*^RC/RC^ mice. Human ADPKD kidney tissues were provided by the NIDDK PKD-Research Resource Consortium and the Kansas PKD Research and Translational Core Center-U54 DK126126.

