## Supplemental figure and Table 1 and will be used for the link to the file preprint site for "Pirfenidone treatment attenuates fibrosis in autosomal dominant polycystic kidney disease"

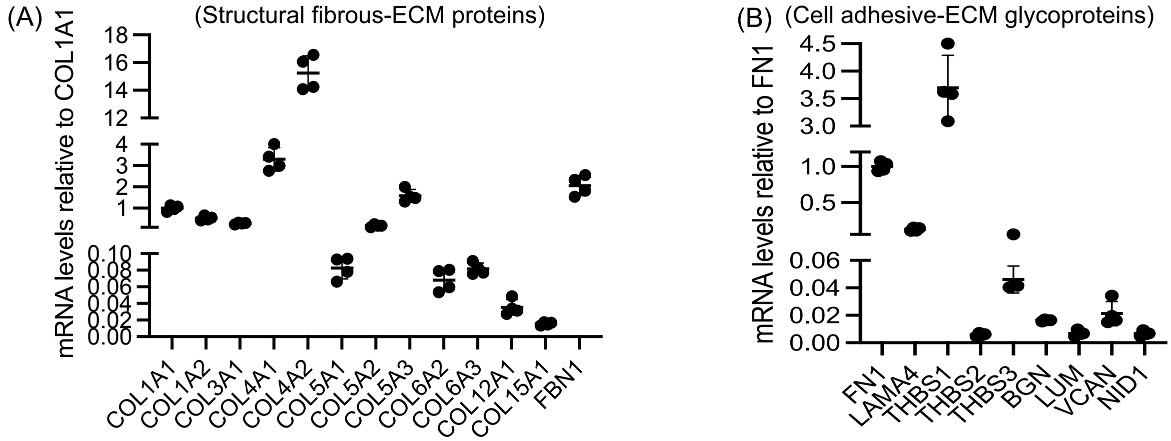


**Supplemental Figure 1:** (A) In primary culture human ADPKD myofibroblasts, relative gene expression of structural fibrous ECM proteins, (B) cell adhesive ECM-glycoproteins.

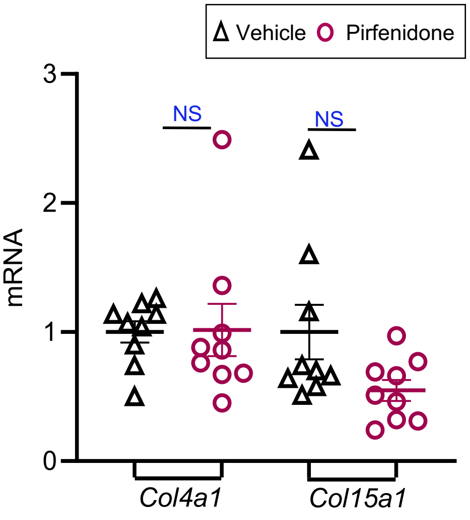


**Supplemental Figure 2:** mRNA levels relative to 18S for Col4a1 and Col15a1 in RC/RC mouse renal tissues.

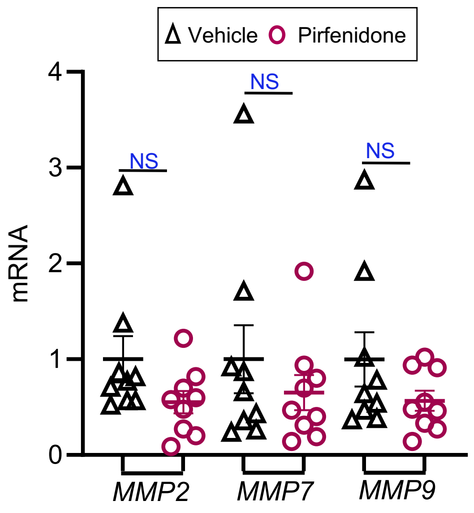


**Supplemental Figure 3:** mRNA levels relative to 18S for MMPs in RC/RC mice renal tissues.

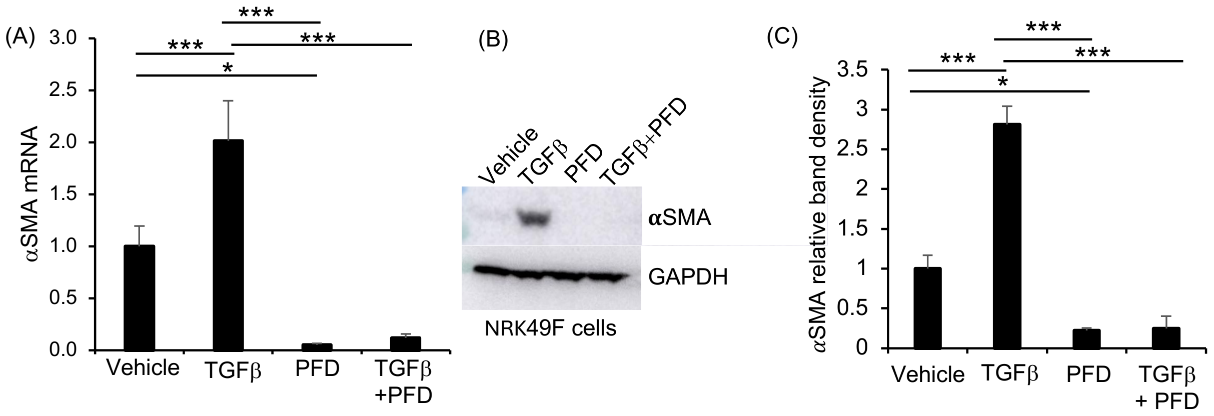


**Supplemental Figure 4:** (A) mRNA levels relative to 18S for αSMA in NRK-49F cells treated with TGFβ and pirfenidone for 24h. (B) Western blot analysis of NRK-49F cells treated with TGFβ (2ng/ml) and pirfenidone (0.5mg/ml) for 24h.(C) quantification of band density. *P<0.05, **P<0.01, ***P<0.001 by two-tailed unpaired t-test with Welch's correction.

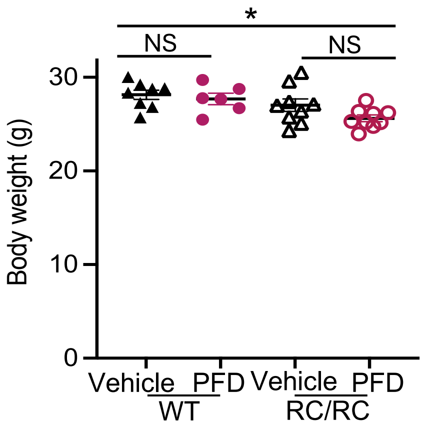


**Supplemental Figure 5:** Body weight of mice treated with vehicle or PFD.

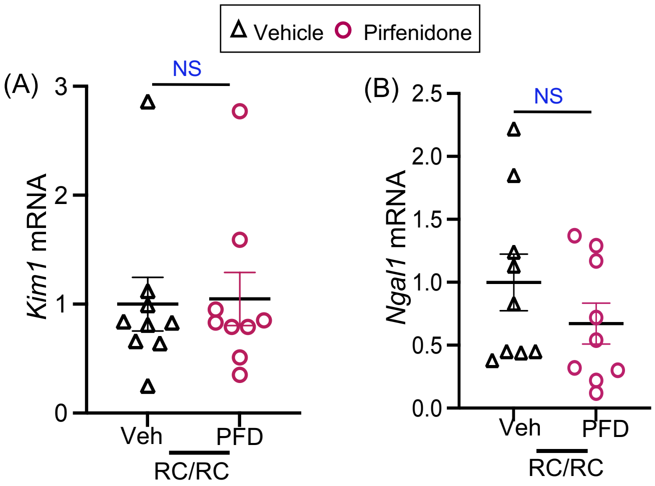


**Supplemental Figure 6:** (A) mRNA levels relative to 18S for Kim1 and (B) Ngal in mouse kidey tissue.

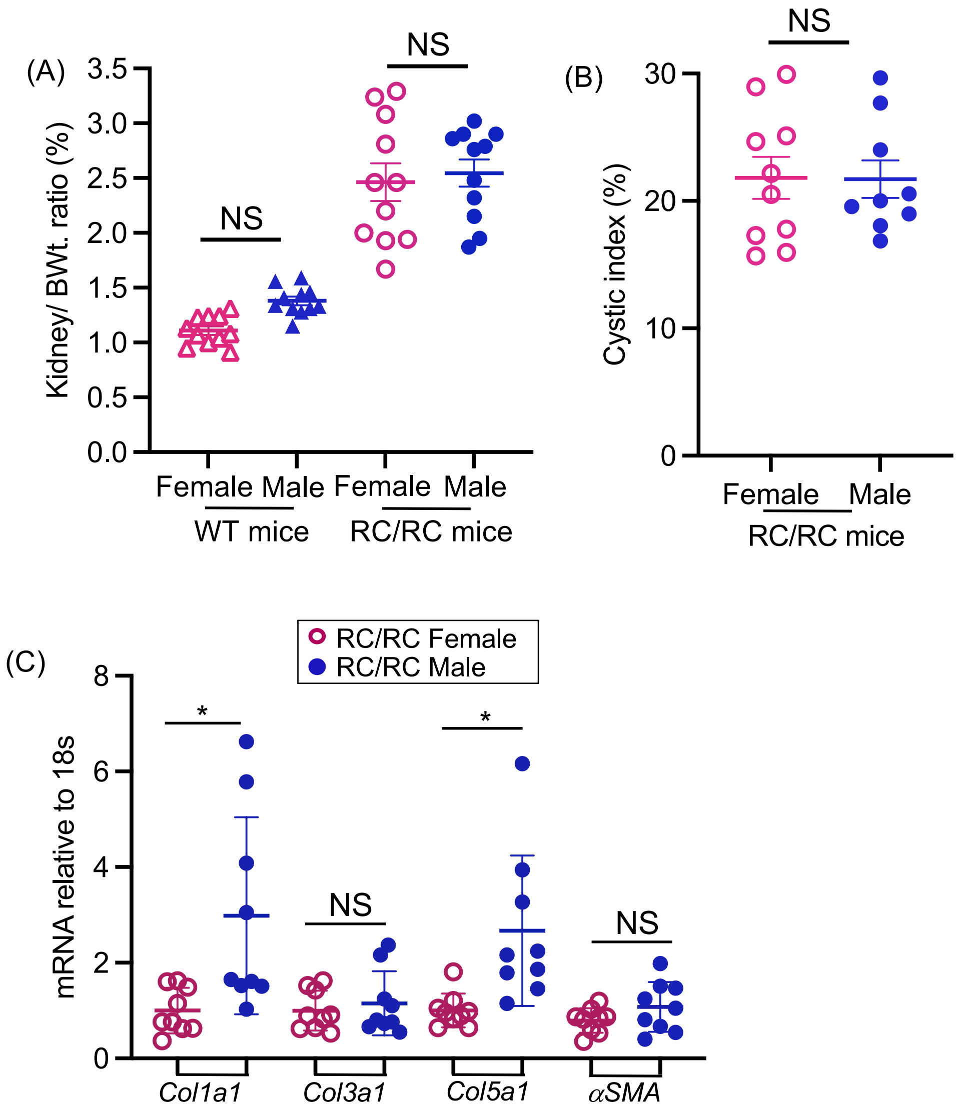


**Supplemental Figure 7:** Male and female RC/RC mice at 6 months of age. (A) Kidney to body weight ratio, (B) Cyst index. (C) mRNA levels relative to 18S for *Col1a, Col3a1, Col5a1* and *αSMA.* Data are normalized to female mice and expressed as fold change. $P<0.05$ by two-tailed unpaired t-test with Welch's correction.

**Supplemental Table-1**

**Human primer sequence used for QRT-PCR**

| | Gene Name | Primer Forward (F) and Reverse (R) |
|---|---|---|
| 1. | MMP-2 | F-AGCGAGTGGATGCCGCCTTTAA<br>R-CATTCCAGGCATCTGCGATGAG |
| 2. | MMP-7 | F-TCGGAGGAGATGCTCACTTCGA<br>R- GGATCAGAGGAATGTCCCATACC |
| 3. | LAMA-4 | F-GAGATGACTCTCTGCTGGACCT<br>R-AGTTCCAGGCAGCCAACAAAGC |
| 4. | TIMP-1 | F-GGAGAGTGTCTGCGGATACTTC<br>R-GCAGGTAGTGATGTGCAAGAGTC |
| 5. | TIMP-2 | F-ACCCTCTGTGACTTCATCGTGC<br>R-GGAGATGTAGCACGGGATCATG |
| 6. | TIMP-3 | F-TACCGAGGCTTCACCAAGATGC<br>R-CATCTTGCCATCATAGACGCGAC |
| 8. | BGN | F-TTGAACCTGGAGCCTTCGATGG<br>R-TTGGAGTAGCGAAGCAGGTCCT |
| 9. | LUM | F-AACATACCAACTGTCAATGAAAACC<br>R-TGCCATCCAAACGCAAATGCTTG |
| 10. | FBN1 | F-GGATACACAGGTGATGGCTTCAC<br>R-GTCGCATTCACAGCGGTATCCT |
| 11. | VCAN | F-TTGGACCTCAGGCGCTTTCTAC<br>R-GGATGACCAATTACACTCAAATCAC |
| 12. | NID1 | F-ACATTGAGCCCTACACGGAGCT<br>R-GCCACTGGTAAGTGTAGATGCG |
| 13. | ADAM12 | F-ATGGCATCTGCCAGACTCACGA<br>R-GGAACTCTTCGAGACTTTGCCAC |
| 14. | ADAM17 | F-AACAGCGACTGCACGTTGAAGG<br>R-CTGTGCAGTAGGACACGCCTTT |
| 15. | ADAM19 | F-CGAGAAGGTGAATGTGGCAGGA<br>R-AGCTCTGACACTGGATCTTCCC |
| 16. | THBS1 | F-GCTGGAAATGTGGTGCTTGTCC<br>R-CTCCATTGTGGTTGAAGCAGGC |
| 17. | THBS2 | F-CAGTCTGAGCAAGTGTGACACC<br>R-TTGCAGAGACGGATGCGTGTGA |
| 18. | THBS3 | F-GGAAGGAGATGCCTGTGACAAC<br>R-GGTAGGATTGCTCATTTCAGGGC |

| | | |
|---|---|---|
| 19. | SPARC | F-TGCCTGATGAGACAGAGGTGGT<br>R-CTTCGGTTTCCTCTGCACCATC |
| 20. | FBLN1 | F-GTGGCTACCATCTCAACGAGGA<br>R-CTTGCATTCGCAGCGGAAACTG |
| 21. | FBLN5 | F-CTGCTGGATGACAACCGAAGCT<br>R-GATAAGGCTCCTCACAGCGGAT |
| 22. | CCN1 | F-GGAAAAGGCAGCTCACTGAAGC<br>R-GGAGATACCAGTTCCACAGGTC |
| 23. | CCN2 | F-CTTGCGAAGCTGACCTGGAAGA<br>R-CCGTCGGTACATACTCCACAGA |
| 24. | COL1a1 | F-TGACGTGATCTGTGACGAGAC<br>R-GGTTTCTTGGTCGGTGGGT |
| 25. | COL1a2 | F-CCTGGTGCTAAAGGAGAAAGAGG<br>R-ATCACCACGACTTCCAGCAGGA |
| 26. | COL3a1 | F- TGG TCT GCA AGG AAT GCC TGGA<br>R- TCT TTC CCT GGG ACA CCA TCAG |
| 27. | COL4a1 | F-TGTTGACGGCTTACCTGGAGAC<br>R-GGTAGACCAACTCCAGGCTCTC |
| 28. | COL4a2 | F-GGATAACAGGCGTGACTGGAGT<br>R-CTTTGCCACCAGGCAGTCCAAT |
| 29. | COL5a1 | F- GGAGATGATGGTCCCAAAGGCA<br>R- CCATCATCTCCTTTGTCACCAGG |
| 30. | COL5a2 | F- CAGGCTCCATAGGAATCAGAGG<br>R- CCAGCATTTCCTGCTTCTCCAG |
| 31. | COL5a3 | F-GAGAGGAGAACTGGGCTTCCAA<br>R-TAGAGGTCCCACTTCTCCTGTC |
| 32. | COL6a2 | F-CGTGGAGACTCAGGACAGCCA<br>R-CCTTTCAAGCCAAAGTCGCCTC |
| 33. | COL6a3 | F-CCTGGTGTAACTGATGCTGCCA<br>R-AAGATGGCGTCCACCTTGGACT |
| 34. | COL12a1 | F-CAGTGCCTGTAGTCAGCCTGAA<br>R-GGTCTTGTTGGCTCTGTGTCCT |
| 35. | COL15a1 | F- GGTGACACTGGTTTACCTGGCT<br>R-GCCTTTCCAGAGGAATGTCCTC |
| 36. | POSTN | F- TGCCCAGCAGTTTTGCCCAT<br>R- CGTTGCTCTCCAAACCTCTA |
| 37. | SERPINE1 | F- CTCATCAGCCACTGGAAAGGCA<br>R- GACTCGTGAAGTCAGCCTGAAAC |
| 38. | FBLN1 | F- GTGGCTACCATCTCAACGAGGA<br>R- CTTGCATTCGCAGCGGAAACTG |
| 39. | FBLN5 | F- CTGCTGGATGACAACCGAAGCT<br>R- GATAAGGCTCCTCACAGCGGAT |
| 40. | 18s | F-ACCGCGGTTCTATTTTGTTG<br>R- CCCTCTTAATCATGGCCTCA |

### Mouse primer sequence used for QRT-PCR

| | Gene Name | Primer Forward (F) and Reverse (R) |
|---|---|---|
| 1. | Adam12 | F-TGCTACAACGGCATCTGCCAGA<br>R-GCTCTTGGAGTCTTTGCCACAG |
| 2. | Adam17 | F-TGTGAGCGGTGACCACGAGAAT<br>R-TTCATCCACCCTGGAGTTGCCA |
| 3. | Adam19 | F-GTGCCTCACTTACCAGGAACAG<br>R-GGACTGCACTTCCTGTATTGGC |
| 4. | Thbs1 | F-GGTAGCTGGAAATGTGGTGCGT<br>R-GCACCGATGTTCTCCGTTGTGA |
| 5. | Thbs2 | F-GTATGGAGGGAAGGACTGTGTC<br>R-ACTTGGCTCCAGGAAAACACGG |
| 6. | SPARC | F-CACCTGGACTACATCGGACCAT<br>R-CTGCTTCTCAGTGAGGAGGTTG |
| 7. | Lama4 | F-CAGTTTGTCCTCTACCTCGGAAG<br>R-CTCACAGGCTTGGAATCCAGGA |
| 8. | Bgn | F-TGAACCAGGAGCCTTTGATGGC<br>R-GTCCTCCAACTCAATAGCCTGG |
| 9. | Vcan | F-GGACCAAGTTCCACCCTGACAT<br>R-CTTCACTGCAAGGTTCCTCTTCT |
| 10. | Col1a1 | F - AGACATGTTCAGCTTTGTGGAC<br>R - GCA GCT GAC TTC AGG GATG |
| 11. | Col1a2 | F-TTCTGTGGGTCCTGCTGGGAAA<br>R-TTGTCACCTCGGATGCCTTGAG |
| 12. | Col3a1 | F - TCC CCT GGA ATC TGT GAA TC<br>R - TGA GTC GAA TTG GGG AGA AT |
| 13. | Col4a1 | F-CGGGTGTGAAAAGACCTATCGG<br>R-CTGGCATTCCTCTGACGCCTTT |
| 14. | Col4a2 | F-CGGGTGTGAAAAGACCTATCGG<br>R-CTGGCATTCCTCTGACGCCTTT |
| 15. | Col5a1 | F-AGATGGCATCCGAGGTCTGAAG<br>R-GACCTTCAGGACCATCTTCTCC |
| 16. | Col5a3 | F-GGCAAAGATGGTATTCCAGGACC<br>R-TGCTTCCTTTGTGACCAGGCATC |
| 17. | Col5a2 | F-GTGGCATAGGAGAGAAAGGTGC<br>R-GCCAACTAAGCCTCTAGGACCA |
| 18. | Col6a1 | F-GACACCTCTCAGTGTGCTCTGT<br>R-GCGATAAGCCTTGGCAGGAAATG |
| 19. | Col6a3 | F-CCTGGTGTAACTGATGCTGCCA<br>R-AAGATGGCGTCCACCTTGGACT |
| 20. | Col8a1 | F-GGAAATCCCACCTGTGCCAAGA<br>R-TCCTCTTGGTCCAGGTTCTCCA |
| 21. | Col12a1 | F-CAGCACCATGAATGTCGTCTGG<br>R-GGTCTTTGAGGATAGTGCTGGC |
| 22. | Col15a1 | F-ACACCCACAGTGACTCCCAAGA<br>R-TCCTCATTGCCCACGATGTCTC |

| | | |
|---|---|---|
| 23. | Col18a1 | F-GTGACACTGGACCTCAAGGCTT<br>R-TTGTCTGAAGGAGGGTCCTGGT |
| 24. | CCN1 | F-GTGAAGTGCGTCCTTGTGGACA<br>R-CTTGACACTGGAGCATCCTGCA |
| 25. | CCN2 | F - GTGCCAGAACGCACACTG<br>R - CCCCGGTTACACTCCAAA |
| 26. | αSMA | F - TCAGGGAGTAATGGTTGGAATG<br>R - GGTGATGATGCCGTGTTCTA |
| 27. | 18s | F - GTAACCCGTTGAACCCGA<br>R - CCATCCAATCGGTAGTAGCG |
| 28. | Timp1 | F - CAAGGATGGACTCCTGGCACAT<br>R - TACTCGCCATCAGCGTTCCCAT |
| 29. | Timp2 | F - AGCCAAAGCAGTGAGCGAGAAG<br>R - GCCGTGTAGATAAACTCGATGTC |
| 30. | Timp3 | F- AGGATGCCTTCTGCAACTCCGA<br>R- GTGTAGACCAGAGTGCCAAAGG |
| 31. | Pai-1 | F - GAGGTGGAAAGAGCCAGATTTA<br>R- CCACTGAAGTAGAGGGCATTC |
| 32. | Mmp2 | F - CAAGGATGGACTCCTGGCACAT<br>R - TACTCGCCATCAColGCGTTCCCAT |
| 33. | Mmp7 | F - AGGTGTGGAGTGCCAGATGTTG<br>R - CCACTACGATCCGAGGTAAGTC |
| 34. | Mmp9 | F - GCTGACTACGATAAGGACGGCA<br>R - TAGTGGTGCAGGCAGAGTAGGA |
| 35. | Hai1 | F- CTTCGTGAGGAAGAGTGCATGC<br>R-TCACACTCCAGGAAGCCATCGA |
| 36. | Areg | F CAGAAGAATGGAAGAGTCAG<br>R CAGATATGCAGGGAGTCACC |
| 37. | Opn | F TGAGAGCAATGAGCATTCCGATG<br>R CAGGGAGTTTCCATGAAGCCAC |
| 38. | He4 | F - AGGTCAAGTCTCCACGAAGCCA<br>R - AGAACACTGGCTGTCCACCTGA |
| 39. | Ngal | F - ATGTCACCTCCATCCTGGTCAG<br>R - GCCACTTGCACATTGTAGCTCTG |
| 40. | Kim1 | F-AAACCAGAGATTCCCACACG<br>R-GTCGTGGGTCTTCCTGTAGC |
| 41. | Fn1 | F -ATGTGGACCCCTCCTGATAGT<br>R -GCCCAGTGATTTCAGCAAAGG |
| 42. | Icam1 | F -CTTCCAGCTACCATCCCAAA<br>R -CTTCAGAGGCAGGAAACAGG |
| 43. | IL-6 | F- CTTCCATCCAGTTGCCTTCT<br>R- CTCCGACTTGTGAAGTGGTATAG |
| 44. | IL-10 | F-TTTGAATTCCCTGGGTGAGAA<br>R-ACAGGGGAGAAATCGATGACA |
| 45. | Tnf | F-ACCCTCACACTCAGATCATCTTC<br>R-TGGTGGTTTGCTACGACGT |
| 46. | Ifn | F -GGCCATCAGCAACAACATAAGCGT<br>R-TGGGTTGTTGACCTCAAACTTGGC |
| 47. | Ccl2 | F -CTCGGACTGTGATGCCTTAAT<br>R- TGGATCCACACCTTGCATTTA |
| 48. | Ccl3 | F-GAAGATTCCACGCCAATTCATC<br>R-GATCTGCCGGTTTCTCTTAGTC |

| | | |
|---|---|---|
| 49. | IL1b | F -TTGACGGACCCCAAAAGAT<br>R - GAAGCTGGATGCTCTCATCTG |
| 50. | Postn | F-CAAAGCACACAGTTACCTTTCCAGGG<br>R-GCAGGAAACCCACATTGCATGAGA |
